# Programmable Allosteric DNAzyme Coupled with CRISPR/Cas12a System for Multiplexed and Sensitive Detection of Extracellular Vesicle Derived MicroRNAs

**DOI:** 10.64898/2026.08.07.743433

**Authors:** Xiang-Lan He, Le Wang, Cong Zhang, Meng-Meng Pan, Ying Ma, Jia-Qi Du, Ling-Jiao Yang, Ming Wang, Ming Jiang, Xu Yu, Li Xu

## Abstract

Extracellular vesicle (EV)-derived microRNAs serve as important biomarkers for cancer diagnosis, yet their accurate detection remains limited by insufficient control of nucleic acid recognition and signal activation. Here, we identified a previously unrecognized feature of CRISPR/Cas12a, in which incorporation of ribonucleotides into single stranded DNA targets modulates Cas12a activation efficiency, revealing a hybrid DNA/RNA-dependent regulation of Cas12a activity. Leveraging this mechanism, we established a programmable detection strategy that enables sequence dependent tuning of Cas12a activation without the need for target amplification. By coupling DNAzyme mediated cleavage with Cas12a *trans*-cleavage, a cascade signal amplification system was established, enabling highly sensitive and selective detection of miRNAs. To facilitate clinical applications, an EV-based sample processing strategy was integrated to simplify isolation of EV associated miRNAs and allow direct miRNA detection without conventional RNA extraction. The resulting platform demonstrated robust discrimination of multiple miRNA targets in clinical cohorts and supported accurate classification of cancer subtypes according to expression signatures. By integrating machine learning analysis, the system accurate distinguished breast cancer (BC) patients from healthy donors (HD), as well as triple-negative breast cancer (TNBC) from BC. This study provides a mechanism-guided strategy for programmable CRISPR-based nucleic acid detection in complex biological samples.

## INTRODUCTION

MicroRNAs (miRNAs) function as central regulators of gene expression and have emerged as important biomarkers for cancer diagnosis due to their stability in circulation and their ability to reflect dynamic changes in cellular states ^1–3^. However, accurate detection of low-abundance miRNAs in complex biological samples remains challenging ^4–5^. This difficulty arises from not only their limited abundance but also insufficient control over nucleic acid recognition and signal activation in current molecular detection systems. Although amplification strategies can improve sensitivity, they often rely on multiple enzymes or complex reaction networks, which increase system complexity and reduce robustness ^6–8^. These limitations highlight the need for strategies that enable precise and programmable control of nucleic acid-triggered signal generation.

Recently, CRISPR-based detection has emerged in molecular diagnosis owing to its high sensitivity, sequence-targeting specificity, and rapid response ^9–11^. In particular, CRISPR/Cas12a system has been widely utilized for nucleic acid and non-nucleic target detection due to its target-activated trans-cleavage activity, which allows signal amplification without extensive reaction cascades ^12–15^. Upon binding to a complementary activator, Cas12a undergoes conformational activation and subsequently cleaves surrounding single-stranded DNA reporters ^16–19^. Despite its utility, the activation of CRISPR-Cas12a is typically governed by strict sequence requirements or crRNA engineering strategies. For example, activation can be regulated through ultraviolet light or target triggered release of caged crRNA ^20–22^, through the use of truncated crRNA or split crRNA designs that maintain partial complementarity to the target ^23^, or through small molecule modulators such as heparin sodium that disrupt the binding between Cas12a and crRNA ^24^. Nevertheless, its responsiveness to subtle variations in target composition remains poorly understood ^25–26^. Meanwhile, how chemical modifications of DNA activators or hybrid nucleic acid features influence Cas12a activation ^27–28^, especially in split target configurations, has not been systematically investigated. This gap limits the rational design of CRISPR-based detection systems with tunable activation behavior.

To overcome the limitations of amplification-dependent detection strategies, deoxyribozymes (DNAzymes) have been widely explored as enzyme-free catalytic modules capable of signal amplification through substrate recycling ^29–31^. DNAzymes are *in vitro* selected DNA molecules that possess various enzyme-like catalytic activities to catalyze multiple chemical reactions, such as repeatedly cleave substrates under target activation ^30, 32^, RNA ligation, and DNA dephosphorylation ^31,33^. They provide an attractive route for enzyme-free signal amplification through catalytic turnover and simple nucleic acid architectures. Integration of DNAzymes with CRISPR systems has enabled enhanced detection sensitivity while reduced reliance on protein enzymes. However, most existing DNAzyme-Cas12a strategies for nucleic acid targets largely treat DNAzymes as auxiliary amplification modules and do not address the fundamental challenge of regulating CRISPR activation at the molecular level or simplifying the overall nucleic acid design ^34–37^. As a result, achieving both high sensitivity and selective discrimination of the target within a simplified system remains difficult.

Here, we identify a previously unrecognized feature of CRISPR/Cas12a activation in which the incorporation of ribonucleotides into single-stranded DNA activators significantly modulates trans-cleavage efficiency of Cas12a in a split target configuration. This finding reveals a hybrid DNA/RNA-dependent regulatory mechanism of Cas12a activation and suggests that the composition can serve as an additional regulatory dimension for programmable control of Cas12a function. By exploiting this property, we established a strategy to convert weakly- or non-activating nucleic acid targets into programmable CRISPR triggers through rational sequence design.

Based on this mechanism, we developed a **<u>D</u>**NAzyme-mediated target-**i**nitiated **<u>s</u>**equential **<u>c</u>**leavage **<u>s</u>**trategy **(DISCs)**, in which target recognition is converted into controlled generation of Cas12a activators through enzyme-free catalytic cascades. This system enables programmable tuning of CRISPR activation without requiring external amplification, while maintaining high sensitivity and sequence specificity through rational nucleic acid design. This design enables enzyme-free amplification signal generation while maintains high sensitivity and sequence selectivity, and significantly simplifies the nucleic acid design by requiring only a minimal set of DNA/RNA components. The modular nature of this system further allows flexible adaptation to different miRNA targets through simple sequence replacement. To enable application in clinically relevant settings, an extracellular vesicle (EV)-enrichment strategy was further incorporated to facilitate efficient isolation of EV-associated miRNAs from complex biological samples. This strategy combines size-recognition using bowl-like TiO_2_ nanospheres with membrane-interacting peptides that would be inserted into lipid bilayers, enabling efficient and unbiased capture of EVs ^38^. Following thermal lysis, released miRNAs were directly introduced into the **DISCs** system without the need for conventional RNA extraction procedures, thereby streamlining sample processing. The **DISCs** system enabled quantification of miRNAs at femtomolar sensitivity.

To demonstrate applicability in complex biological samples, this strategy was applied to the analysis of EV-associated miRNAs in clinical samples. Using this approach, multiple miRNAs can be profiled, supporting the discrimination of cancer subtypes based on expression patterns. By integrating machine learning to process data, the system achieved highly accurate classification in a cohort of 65 clinical samples, distinguishing breast cancer (BC) from healthy donors (HD), as well as triple-negative breast cancer (TNBC) from BC. This work establishes a mechanism-guided framework for regulating CRISPR activity and provides a versatile strategy for nucleic acid detection in complex biological systems.

## RESULTS AND DISCUSSION

### Inhibitory effect of RNA-containing ssDNA targets on Cas12a activity

In a canonical Cas12a reaction, the Cas12a associates with crRNA to form a functional ribonucleoprotein (RNP) that recognizes double stranded or single stranded DNA targets (dsDNA/ssDNA) containing sequences complementary to the crRNA spacer region. Upon target recognition, the Cas12a RNP undergoes a conformational alteration that initiates both cis-cleavage of the target and trans-cleavage of surrounding ssDNA substrates. The activated RNP repeatedly cleaves fluorophore quencher-labeled ssDNA reporters (F-Q reporters), generating amplified fluorescence signals without sequence specificity ^16, 39–40^. This collateral cleavage activity constitutes the central signal amplification mechanism of Cas12a based biosensors and has been widely exploited for molecular diagnostics, as illustrated in **Fig. 1A**. According to this mechanism, formation and stabilization of the Cas12a/crRNA/target ternary complex determined whether the trans-cleavage activity can be efficiently triggered or not. Minor perturbations in the activator sequence may disrupt the stability of the ternary complex and thereby weaken or abolish activation of collateral cleavage.

To systematically examine the tolerance of Cas12a toward sequence variations in ssDNA activators, we investigated the trans-cleavage response of Cas12a toward a series of ssDNA activators with different sequence features. As shown in **Fig. 1B** and **Fig. 1D**, the fully complementary DNA activator generated a strong fluorescence signal, confirming that the trans-cleavage activity of Cas12a was efficiently initiated. Interestingly, an ssDNA activator containing two mismatched bases (DNA-mis) still produced a pronounced fluorescence signal, corresponding to an activation efficiency of approximately 59.44% ± 2.07%. This observation indicates that Cas12a exhibits substantial tolerance toward limited base mismatches in ssDNA. Such mismatch tolerance may arise from the flexible recognition process between the crRNA spacer region and the target sequence during R loop formation as reported previously ^41–42^.

Recent reports suggest that split-activator combinations can partially activate Cas12a trans-cleavage activity when the combined length of two fragments (ssDNA/ssDNA or ssDNA/RNA) reaches or exceeds 20 nt nucleotides ^43–45^. To examine this phenomenon, two pairs of split ssDNA fragments were designed. One pair contained fragment with 10 nt complementary to the crRNA-DR spacer region, denoted as **D-1** + **D-2**, while the other pair contained fragments with 9 nt complementary nucleotides, donated as **D-3** + **D-4** (**Fig. 1B**). As presented in **Fig. 1B** and **Fig. 1D**, the **D-3** + **D-4** combination yielded fluorescence signals comparable to the negative control, indicating negligible activation of Cas12a. In contrast, **D-1** + **D-2** combination generated only a weak fluorescence signal, corresponding to a residual activation efficiency of 7.93% ±1.18% relative to the positive control. These findings demonstrate that the structural integrity of the ssDNA activators strongly influences Cas12a activation, and fragmentation of the activator substantially compromises the ability to stabilize the ternary complex.

Previous studies have established that DNA serves as the predominant activator of Cas12a trans-cleavage activity, whereas RNA targets are largely refractory to triggering this cleavage activity ^22, 46^. Based on this property, we sought to investigate whether incorporation of ribonucleotides into ssDNA activators could modulate the DNA target-mediated activation of Cas12a. Accordingly, a DNA/RNA chimeric activator was therefore designed by introducing two ribonucleotides at positions 9 and 10 downstream of the seed region. As shown in **Fig. 1C** and **Fig. 1E**, the intact DNA/RNA activator retained robust trans-cleavage activity and generated strong fluorescence signals. Although the FL intensity was slightly lower than that obtained with the pure DNA activator, the activation efficiency still reached approximately 90.95% ±1.94%. These results indicate that limited incorporation of ribonucleotides does not significantly disrupt target recognition, and Cas12a can tolerate ribonucleotides at positions adjacent to the seed region.

Interestingly, when the two ribonucleotides were replaced with mismatched ribonucleotides, the trans-cleavage activation efficiency of Cas12a decreased dramatically to 12.69% ± 1.12%, indicating a substantial reduction compared with DNA-mis. Furthermore, splitting the DNA/RNA activator into two chimeric fragments (**D/R-1** + **D/R-2**) failed to induce detectable Cas12a activation. These findings suggest that the introduction of ribonucleotides enhances the sensitivity of the Cas12a system to both base mismatches and structural disruption of the activator. The presence of ribonucleotides near the PAM “seed” region may alter local hybridization stability or conformational dynamics of the crRNA target duplex, thereby reducing the tolerance of Cas12a toward imperfect targets.

In summary, these results reveal that although Cas12a can accommodate limited ribonucleotide incorporation within DNA activators, such modification markedly decreases the tolerance of the system toward mismatches and fragmented targets. This property provides a useful strategy for improving sequence discrimination in CRISPR/Cas12a based biosensing systems and lays the foundation for rational design of activator structures with enhanced specificity.

**Figure 1.**
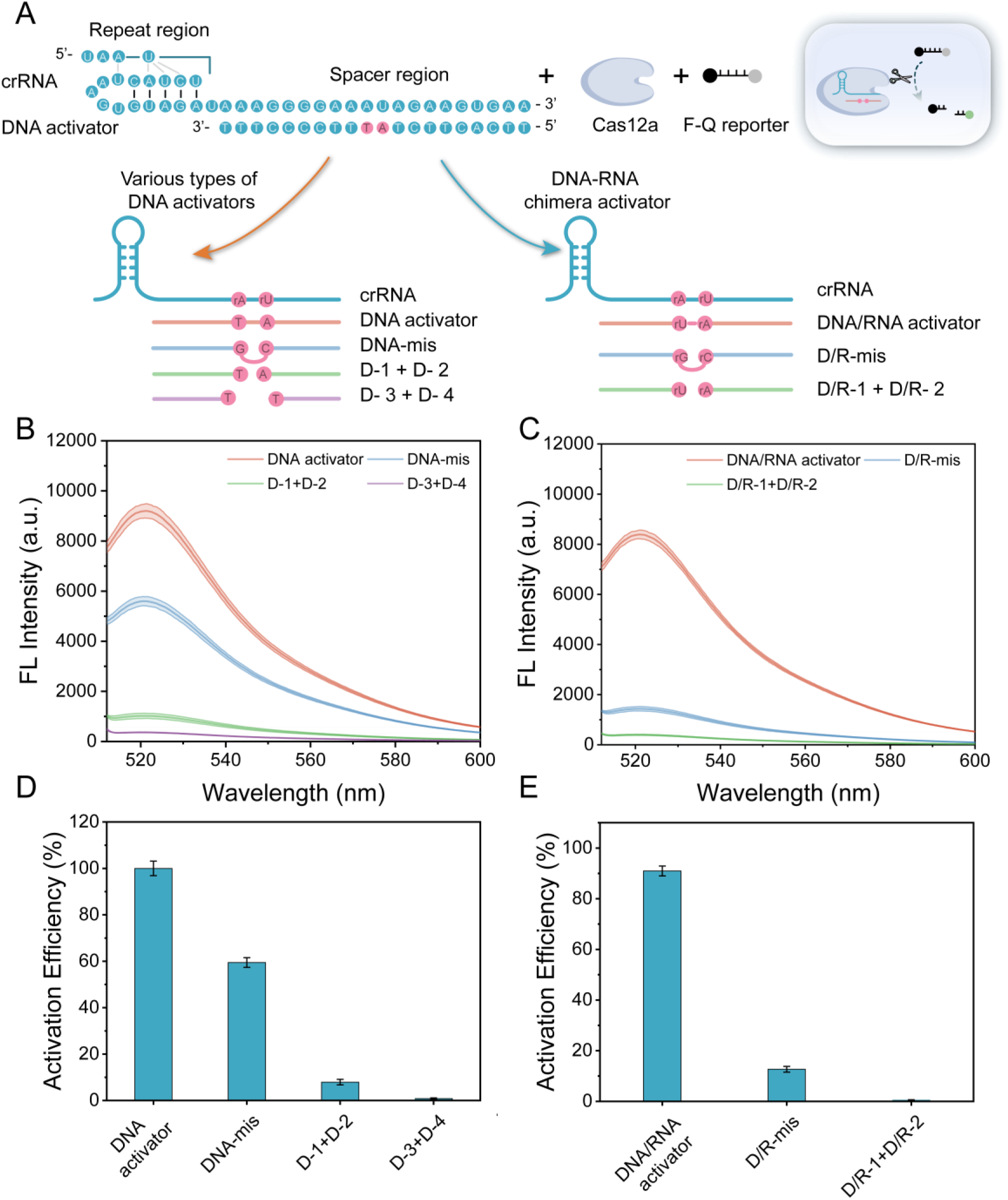
Effects of different types of DNA activators on the activation of cas12a trans-cleavage activity. (A) Illustration of the activation to RNP of the cleavage product. (B) and (C) Fluorescence spectra of the different DNA activators. (D) and (E) Comparison of activation efficiency elicited by different ssDNA activators.

### Design principle of the DISCs system

Inspirated by the above findings, we developed a detection system for miRNA detection, which integrates DNAzyme-mediated target-initiated sequential cleavage with CRISPR/Cas12a cascade signal amplification (**Scheme 1 and Fig. S1**). This design exploits the programmable catalytic activity of DNAzymes together with the collateral trans-cleavage property of Cas12a to construct a sensitive nucleic acid detection platform. DNAzymes recognize their substrates through sequence-specific hybridization and subsequently catalyze the hydrolysis of phosphodiester bonds. Among the reported DNAzymes, the well-characterized 10-23 and 8-17 DNAzymes exhibit efficient endonuclease activity toward ssRNA substrates ^47^. In this study, these catalytic features coupled with the previously observed activation characteristics of Cas12a were rationally utilized to construct a “turn-off” sensing mechanism. To achieve high sensitivity and fast response, the 10-23 DNAzyme was selected as the catalytic core, because of its high catalytic efficiency toward RNA containing substrates, which enables rapid substrate turnover and efficient signal transduction ^33, 48^. To regulate the catalytic activity of the DNAzyme in a target responsive manner, the 5’ end of the 10-23 DNAzyme was extended with a self-folding locking domain that sterically blocks the binding of the DNA/RNA chimeric substrate, denoted as D/R-S. The D/R-S was complementary to the spacer region of crRNA-DR within the Cas12a ribonucleoprotein complex and could therefore activate Cas12a trans-cleavage activity in its intact form. The locking domain was engineered as a loop structure containing a target recognition sequence that served as an allosteric regulatory module controlling the accessibility of the DNAzyme catalytic site.

Based on this design, the target-triggered DNAzyme probe, termed THD, was engineered as a locked hairpin structure comprising three modular sequences, including a target recognition region shown in red, a catalytic core region shown in blue, and scaffold domains that maintained the overall structure framework of the probe (**Fig. S2**). In the presence of target miRNA, the THD probe hybridized with the target miRNA through sequence complementarity, which induced a conformational transition that disrupted the locking structure and exposed the D/R-S-binding domain. This structural rearrangement activated the catalytic function of the DNAzyme, enabling the probe to bind and cleave the D/R-S substrate through iterative catalytic cycles. The DNAzyme-mediated cleavage produced two short DNA/RNA chimeric fragments that were incapable of activating Cas12a trans-cleavage activity, as demonstrated in the preceding experiments. As a result, the cleavage of D/R-S effectively prevented the formation of active Cas12a RNP complexes and led to a reduction in fluorescence signal. In contrast, in the absence of target miRNA, the THD probe remained in its locked conformation and the D/R-S remained intact. The intact D/R-S subsequently activated the Cas12a RNP complex, which triggered non-specifical trans-cleavage of F-Q reporters and produced strong fluorescent signals. Through this cascade regulatory mechanism, the presence of target miRNA suppressed Cas12a activation, whereas its absence resulted in efficient reporter cleavage. The **DISCs** system therefore converted DNAzyme-mediated target recognition into an amplified CRISPR Cas12a fluorescence readout, enabling sensitive detection of miRNA.

**Scheme 1.**
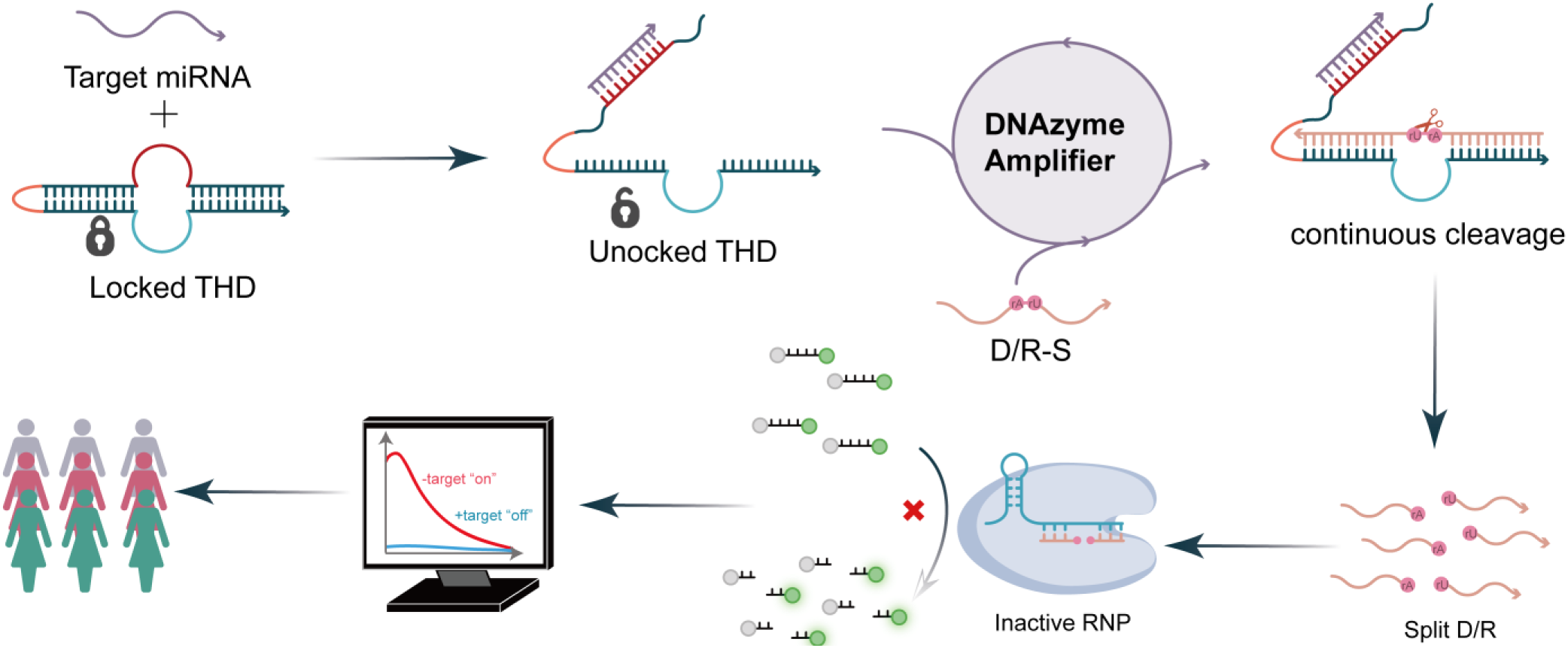
The detection process of the **DISCs** system.

### Feasibility analysis of the DISCs detection process

MicroRNA 21 (miR-21) is one of the most extensively studied oncogenic microRNAs and has been widely reported to participate in tumor proliferation, metastasis, and therapeutic resistance ^49–51^. Elevated expression of miR-21 is frequently associated with poor clinical prognosis across multiple malignancies. Detection of this potential biomarker in EVs might provide valuable information for early cancer diagnosis, mechanistic investigation, and therapeutic monitoring ^19, 52–53^. To establish and optimize the DISCs biosensing platform, a DNA mimic of miR-21 (miD-21), was herein selected as the model target. The native polyacrylamide gel electrophoresis (PAGE) was conducted to assess the target triggered cleavage behavior of the THD. As shown in **Fig. 2A**, lanes 1, 2 and 3 corresponded to the THD probe, D/R-S, and target miD-21, respectively. Upon incubation of THD with the target miD-21, a new band with lower electrophoretic mobility appeared in lanes 5 and 6, indicating the formation of a duplex structure between the THD probe and miD-21 through sequence specific hybridization. It should be noted that, in the presence of target, the D/R-S was cleaved into short fragments by the activated DNAzyme. These fragments were too small to be observed in the gel, leading to the disappearance of the corresponding band in lane 6. In contrast, in the absence of the miD-21, THD probe remained in its locked hairpin conformation, which prevented D/R-S binding and subsequent cleavage. As a result, no additional complexes were formed, and the THD probe maintained its structural stability. To further verify that the cleavage products of D/R-S were incapable of activating Cas12a trans-cleavage activity, additional PAGE analysis was performed. As displayed in **Fig. S3A**, lane 1 corresponded to the intact ssDNA reporter that could be cleaved by activated Cas12a RNP. In lane 2, the disappearance of the ssDNA band indicated efficient cleavage by Cas12a RNP. In contrast, the ssDNA band remained intact in lane 3 when the cleaved D/R-S fragments were introduced, demonstrating that the cleavage products of D/R-S failed to activate the Cas12a RNP. These results confirm that fragmentation of the activator efficiently suppresses the trans-cleavage activity of Cas12a.

We subsequently evaluated the performance of the DISCs system under different experimental conditions. As shown in **Fig. 2B** and **Fig. S3B**, a strong fluorescence signal was observed in the absence of target miD-21 or the THD probe. Under these conditions, the unlocked THD probe failed to cleave D/R-S and the intact D/R-S substrate was able to activate the Cas12a RNP, which triggered trans-cleavage of F-Q report probes and generated pronounced fluorescence signals. In contrast, when all reaction components including target miD-21, THD probe, D/R-S substrate, and Cas12a RNP were simultaneously present, only a negligible fluorescence signal was observed. These observations indicate that the presence of miD-21 activates the THD probe, leading to catalytic cleavage of the D/R-S substrate of the DNAzyme and suppression of Cas12a activation. In summary, the above results demonstrate the feasibility of the DISCs sensing strategy and confirm that target induced DNAzyme activation can effectively regulate Cas12a mediated fluorescence output, thereby enabling sensitive detection of miRNA.

**Figure 2.**
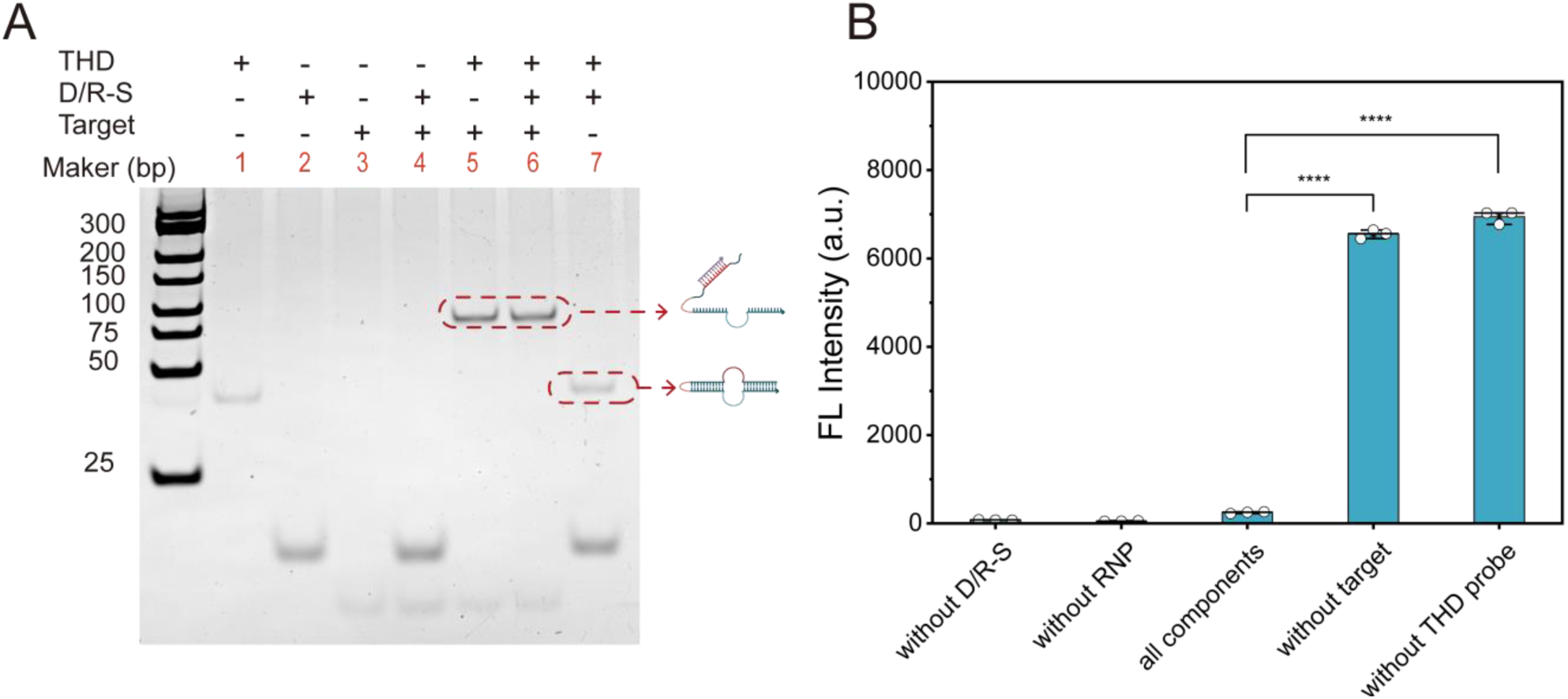
The feasibility of the **DISCs** system. (A) Analysis of the cleavage of the THD probe using the native PAGE. Lane 1: THD; Lane 2: D/R-S; Lane 3: miD-21; Lane 4: D/R-S + miD-21; Lane 5: THD + miD-21; Lane 6: THD + D/R-S + miD-21; Lane 7: THD + D/R-S. (B) FL intensities of the **DISCs** under different conditions. The P-values are represented by asterisks as follows: \**P* < 0.05, \*\**P* < 0.01, \*\*\**P* < 0.001, and \*\*\*\**P* < 0.0001.

### Optimization of the DISCs detection conditions

To maximize the detection performance of the **DISCs** platform, the experimental parameters were systematically optimized. We initially examined the impact of buffer pH and Mg2+ concentration on the sensing performance. As shown in **Fig. 3A**, the F0/F ratio increased progressively with elevated pHs and reached its maximum at pH 9.0, which the F_0_ presents the FL intensity without target and the F presents the FL intensity with target. This trend was consistent with the catalytic mechanism of the 10-23 DNAzyme ^33^. The RNA cleavage catalyzed by the 10-23 DNAzyme involved transesterification reaction, in which the 2’-OH group acted as a nucleophile to attack the adjacent phosphate. Deprotonation of the 2’-OH group by a general base facilitated this nucleophilic reaction, and alkaline conditions therefore promoted catalytic efficiency ^33, 54–55^. The effect of Mg^2+^ concentration was subsequently evaluated, as Mg^2+^ was known to play an essential structural and catalytic role in many DNAzyme reactions. As depicted in **Fig. 3B**, the F_0_/F ratios reached its maximum at a Mg^2+^ concentration of 40 mM, while both lower and higher Mg^2+^ concentrations resulted in diminished signal responses. This phenomenon might be potentially attributed to insufficient structural stabilization at low Mg^2+^ concentrations or excessive ionic strength at high concentrations, both of which could perturb the conformational stability of the THD probe and influence the catalytic efficiency of the DNAzyme.

The concentration of the D/R-S was further optimized. As shown in **Fig. 3C**, the F_0_/F ratio increased progressively with increasing D/R-S concentration and reached a plateau at approximately 80 nM. At this concentration, sufficient substrate molecules were available to efficiently activate Cas12a in the absence of target while still allowing effective signal suppression after DNAzyme mediated cleavage. Subsequently, the concentration of the THD probe was also found to significantly affect the sensing performance. As shown in **Fig. 3D**, increasing the THD probe concentration beyond a certain threshold resulted in a noticeable decrease in the F_0_/F ratio. When the THD concentration reached 200 nM, background fluorescence was substantially reduced, likely due to probe heterogeneity. Because a minor subset of THD probes remains unblocked, increasing the total probe concentration elevates the absolute number of unblocked species, which in turn causes both background and experimental signals to be simultaneously reduced. Therefore, an optimal THD concentration of 80 nM was selected to achieve the highest signal-to-noise ratio.

We further evaluated the effects of the reaction temperature and cleavage time for the THD probe, as well as the incubation temperature for the CRISPR/Cas12a reaction. As shown in **Fig. 3E** and **Fig. S4**, both the DNAzyme-mediated cleavage process and the Cas12a reaction exhibited optimal performance at 25℃. The time dependent cleavage analysis indicated that the catalytic reaction of the THD probe gradually approached a plateau within approximately 3 h (**Fig. 3F**), suggesting that sufficient substrate turnover had been achieved during this period. Consequently, the optimal detection conditions for the **DISCs** detection system were established as follows: buffer pH 9.0 containing 40 mM Mg^2+^, 80 nM D/R-S and 80 nM THD probe, a THD cleavage reaction time of 3 h, and a reaction temperature of 25℃.

**Figure 3.**
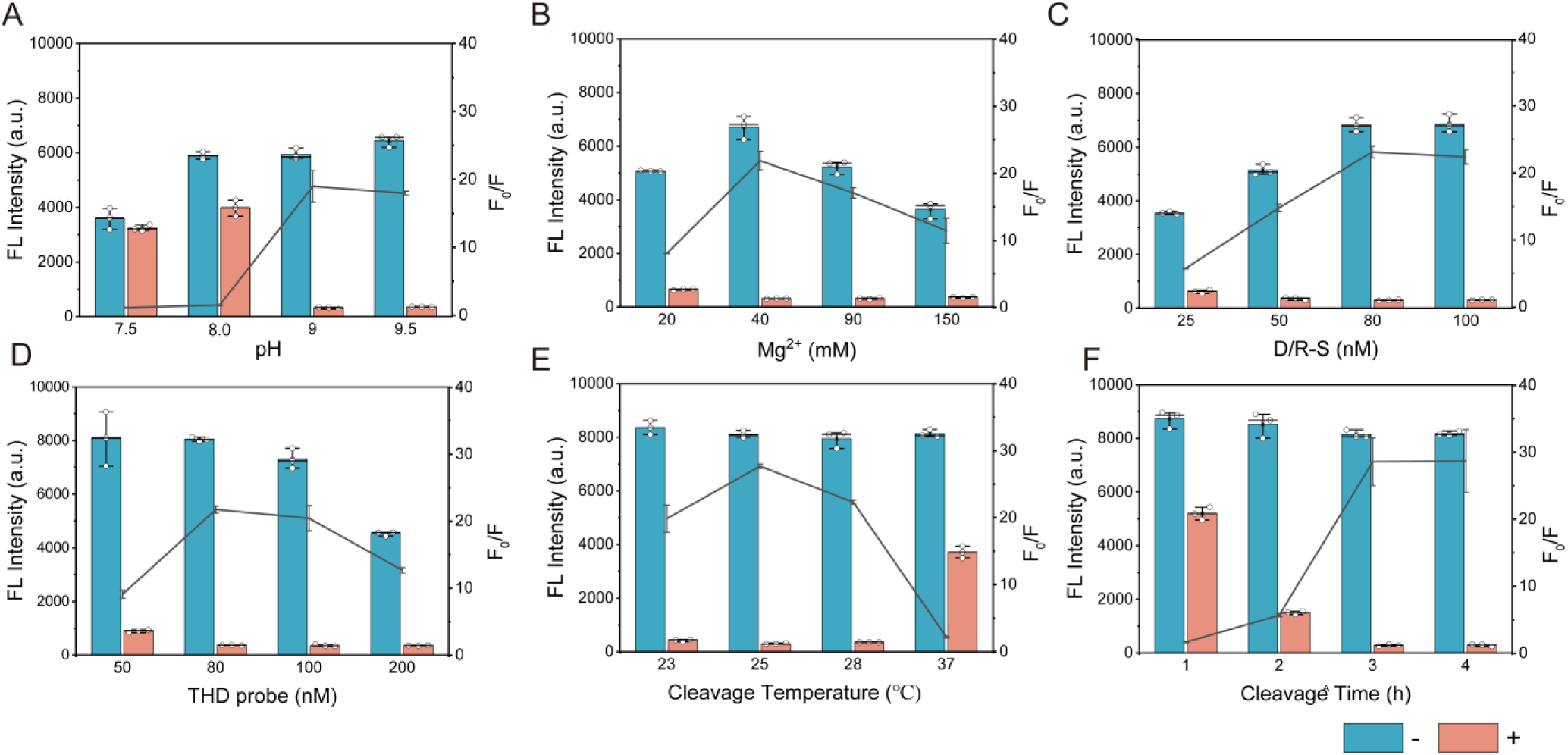
Optimization of **DISCs** system. The effects of key reaction parameters were systematically investigated, including (A) reaction buffer pH, (B) Mg^2+^ concentration, (C) D/R-S concentration, (D) THD concentration, (E) cleavage temperature, and (F) DNAzyme cleavage time.

### Analytical performance of the DISCs assay

Prior to assessing the analytical performance of the DISCs assay, we first compared the activation efficiencies of miR-21 and its DNA mimic miD-21. As shown in **Fig. 4A**, no significant difference in the FL intensity was observed between the two targets (*P* > 0.05), indicating that the THD probe could be activated with comparable efficiency by both miR-21 and miD-21. Encouraged by this, we further investigated the constructed strategy for quantitative analysis of the miR-21. The target miR-21 was serially diluted over a concentration range from 10 fM to 5 nM. As presented in **Fig. 4B** and **Fig. 4C**, the FL intensity exhibited a strong linear relationship with the logarithmic concentration of miR-21, yielding an excellent correlation coefficient of R^2^ = 0.9974. Based on the standard formula 3σ/k, the limit of detection (LOD) was calculated as low as 2.76 fM. This detection sensitivity was comparable to or superior to many previously reported methods (**Table S2**) ^56–60^, highlighting the high amplification efficiency of the DISCs system.

To further demonstrate the versatility of the proposed platform, the detection strategy was extended to other miRNA targets by simply redesigning the target recognition region of the THD probe. As displayed in **Fig. 4D**, although slight differences in FL intensity were observed among different miRNAs detection systems, all assays exhibited high F_0_/F values, indicating that the DISCs platform maintained robust signal responses across different targets. The analytical performance for additional targets, including miR-141 and miR-10b, was subsequently evaluated. Good linear relationships were obtained between the FL intensity and the logarithmic concentrations of both miR-141 and miR-10b over a range from 10 fM to 5 nM, with the corresponding linear equations of y = 8.409 ×10^3^-1.217 ×10^3^ ×log CmiR-141 (R^2^ = 0.9939) and y = 8.989 ×10^3^-1.304 ×10^3^ ×log CmiR-10b (R^2^ = 0.9943), respectively (**Fig. 4E, Fig. S5A, Fig. 4F** and **Fig. S5B**). The calculated LOD were 1.89 fM for miR-141 and 1.10 fM for miR-10b, demonstrating the consistently high sensitivity of the **DISCs** platform.

Because numerous endogenous miRNAs coexisted in clinical samples and might interfere with probe recognition, the specificity of the detection strategy was further investigated through systematic cross reactivity analysis. As depicted in **Fig. 4G** and **Fig. 4H**, only the perfectly matched target miRNAs were able to activate their corresponding THD probes, resulting in a pronounced reduction in FL signals. In contrast, no target miRNAs generated negligible responses, indicating minimal cross interference. Collectively, these findings demonstrate that the DISCs strategy provides a highly sensitive, specific and versatile platform for the universal detection of miRNAs.

**Figure 4.**
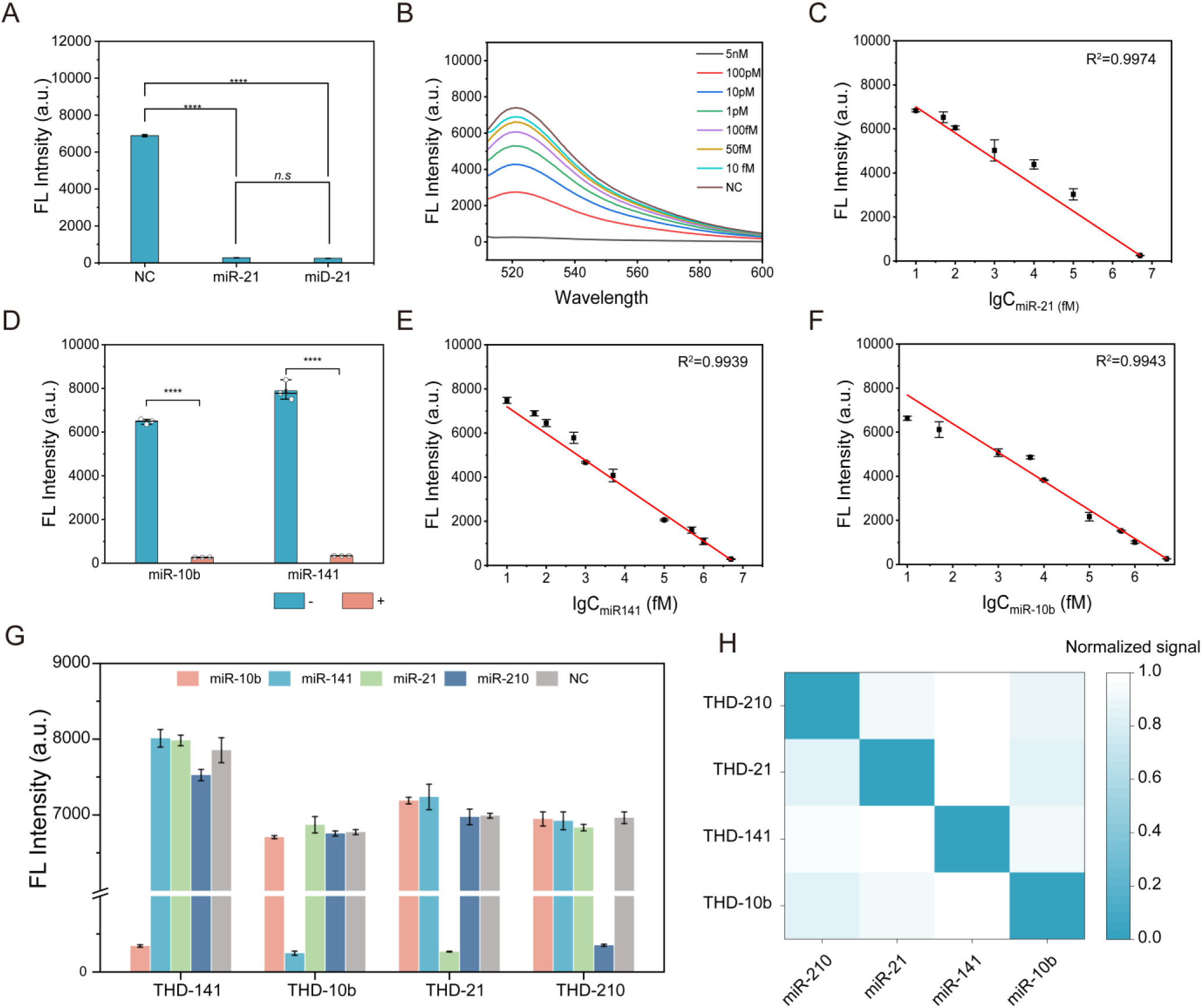
Analytical performance of the DISCs detection assay. (A) Comparative analysis of THD probe activation efficiency in response to miD-21 and miR-21. (B) and (C) Fluorescence response profiles and corresponding linear curves for varying concentrations of miR-21. (D) Validation of sequence-specific activation of target-specific THDs by their cognate miRNAs. Corresponding linear curves for (E) miR-141 and (F) miR-10b. (G) Fluorescence intensity profiles generated using distinct THD probes in the presence of different miRNA species. (H) Heatmap visualization of the normalized signal intensities across different THD probe and miRNAs. Signal normalization was calculated using Z-score transformation. All data are presented as mean values ±SD. Statistical significance is indicated as follows: \**P* < 0.05, \*\**P* < 0.01, \*\*\**P* < 0.001, and \*\*\*\**P* < 0.0001.

### Establishment of an EVs Capture System

Ultracentrifugation (UC) was widely regarded as the gold-standard method for EVs isolation. However, this approach was limited by long processing time, labor intensive procedures, and the requirement for specialized instrumentation and trained personnel ^61–62^. To address these limitations, we developed a rapid EV capture strategy based on concave TiO_2_ nanospheres synthesized via an electrospray approach. As depicted in **Fig. 5A**, the obtained TiO_2_ nanospheres exhibited a distinct bowl-shaped morphology with a concave cavity structure. The average size of the concave cavity structure was approximately 200 nm, which was comparable to the typical size range of EVs. This geometric compatibility enabled effective topological matching between the TiO_2_ nanospheres and EVs, thereby facilitating physical confinement and improving capture efficiency.

To further enhance the specificity and affinity of EV capture, the surface of the TiO_2_ nanospheres was functionalized with **<u>b</u>**iotin modified **<u>a</u>**rtificial **i**nsertion **<u>p</u>**eptide (BAIP). This type of peptide was capable of inserting into lipid bilayers and could therefore selectively anchor onto EV membranes according to our previous report ^38^. Through the combined effects of structure matching between the concave nanospheres and EVs together with the affinity insertion interactions of BAIP, the resulting BAIP-TiO_2_ nanospheres enabled rapid, high-affinity and unbiased capture of EVs even in complex biological environments. After isolation, the cargo molecules encapsulated within the EVs were efficiently released through a simple thermolysis process performed at 90℃ for 10 min. This thermal lysis step efficiently disrupted the vesicular membrane and liberated the enclosed nucleic acids. Compared with conventional RNA extraction procedures, this straightforward approach minimized sample loss and eliminated the need for multiple purification steps or specialized equipment. Consequently, the integration of the BAIP-TiO_2_ capture strategy with rapid thermal release significantly simplified the workflow for downstream detection of EV-derived miRNAs.

### Characterization and Modification of the BAIP-TiO_2_ for EVs Isolation

We systematically characterized the physicochemical properties of the TiO_2_ nanospheres. As shown in **Fig. S6A**, the TiO_2_ nanospheres displayed a distinct bowl-like morphology with unilateral concave indentations. The size of the concave cavity was comparable to that of EVs, ∼ 200 nm, indicating their potential to trap EVs via topological recognition and spatial confinement. X-ray photoelectron spectroscopy (XPS) was utilized to analyze the element composition and chemical states of the TiO_2_ nanosphere. The survey spectrum revealed the presence of four characteristic signals locating at 284.0 eV, 457.8 eV, and 529.1 eV, (**Fig. S6B**), corresponding to C 1s, Ti 2p, and O 1s, respectively. The high-resolution Ti 2p spectrum displayed two distinct peaks at 457.8 eV and 463.4 eV, which could be assigned to Ti 2p3/2 and Ti 2p1/2, respectively, confirming the typical chemical state of Ti^4+^ in TiO_2_ ^63–64^. Simultaneously, the O 1s spectrum exhibited two components centering at 529.0 eV and 531.0 eV, which were attributable to the Ti-O-Ti bonding and -OH group on the TiO_2_ surface, respectively ^65–66^. These results collectively confirme the successful formation of TiO_2_ nanostructures with well-defined chemical composition.

To further improve the capture efficiency, the bowl-like TiO_2_ nanospheres were functionalized with BAIP. The surface modification process was achieved through a stepwise assembly strategy. Initially, BSA-biotin was immobilized onto the TiO_2_ nanospheres surface via nonspecific adsorption ^67^, providing a biotinylated interface for subsequent conjugation. The as-prepared BSA-biotin-TiO_2_ nanospheres were sequentially incubated with SA and BAIP, leading to the formation of BAIP functionalized TiO_2_ nanospheres through strong biotin-SA affinity interactions. The modification conditions were further optimized to ensure efficient surface functionalization. As shown in **Fig. S6C** and **Fig. S6D**, the immobilization efficiency of both SA and BAIP increased with concentration and reached saturation at 100 μg/mL. These findings suggest that successful peptide functionalization of the material surface was achieved under the optimized BAIP and SA conditions, providing a robust platform for subsequent EV membrane anchoring.

### EVs capture by the BAIP-TiO_2_ Nanospheres

The model EVs derived from the MCF-7 cell line were isolated by UC and utilized to evaluate the EV capture capability of BAIP-TiO_2_ nanospheres and to optimize the capture conditions. The model EVs were first characterized by NTA and TEM. As illustrated in **Fig. S7A, Fig. S7B** and **Fig. S7C**, the EVs exhibited a typical cup-shaped morphology with an average diameter of 202 nm, which was consistent with previous results ^68–70^. Western blot analysis was further employed to identify characteristic marker proteins. As shown in **Fig. S7D**, EV specific markers including CD9 and β-actin were detected in EVs, whereas the negative marker GM130 was absent in the EV fraction, confirming the successful isolation and purification of EVs ^71^. Subsequently, the EV capture performance of BAIP-TiO_2_ nanospheres was investigated. As shown in **Fig. 5B**, numerous spherical protrusions were observed on the surface of BAIP-TiO_2_ after incubation with EVs, in sharp contrast to the smooth surface of the bare TiO_2_ nanospheres (**Fig. S6A**). These protrusions were consistent with the size and morphology of EVs, indicating successful capture on the nanosphere surface.

To further verify EV binding, immunofluorescence staining was performed to characterize CD9 protein in the isolated EVs. As displayed in **Fig. 5C, Fig. S7E** and **Fig. S7F**, distinct red fluorescence was observed on BAIP-TiO_2_ which were incubated with EVs under inverted confocal microscopy, whereas only weak fluorescence was detected on the control group. These fundings demonstrate that the BAIP-TiO_2_ nanospheres could efficiently capture EVs. The capture conditions were further optimized to maximize efficiency. As shown in **Fig. 5D** and **Fig. 5E**, an incubation time of 30 min and a BAIP-TiO_2_ nanospheres of 1 mg resulted in a capture efficiency of approximately 80%, indicating rapid and effective EV enrichment under mild conditions. Under the optimal conditions, we further investigated the respective contributions of the bare TiO_2_ nanostructure and BAIP modification. As depicted in **Fig. 5F**, bare bowl-like TiO_2_ nanospheres exhibited an intrinsic EV capture capacity with an efficiency of 33.2% ± 3.0%, which could be attributed to topological recognition and size matching effects. Upon BAIP modification, the capture efficiency increased significantly to 79.4% ± 5.33%. This enhancement arises from the additional membrane insertion and specific interaction provided by BAIP, demonstrating a synergistic effect between nanostructure mediated topology recognition and BAIP peptide mediated affinity recognition. With both the EV capture module and the DISCs detection module established, we integrated a unified workflow for EV-derived miRNA detection, termed **Cap-DISCs**. Mimicking an optical drive, BAIP-TiO_2_ nanospheres “load” target EVs (Cap) while DISCs “read” the encapsulated miRNA. Following capture, EVs were lysed through a simple thermolysis process, releasing encapsulated miRNAs that directly activate the DNAzyme module. This activation initiated a cascade of cleavage events that ultimately suppressed the trans-cleavage activity of Cas12a, yielding a decrease in fluorescence signal (**Fig. 5G**). Collectively, these results demonstrate the successful development of an integrated platform that enabled efficient EV capture and sensitive downstream detection, providing a streamlined strategy for EV-based molecular diagnostics.

**Figure 5.**
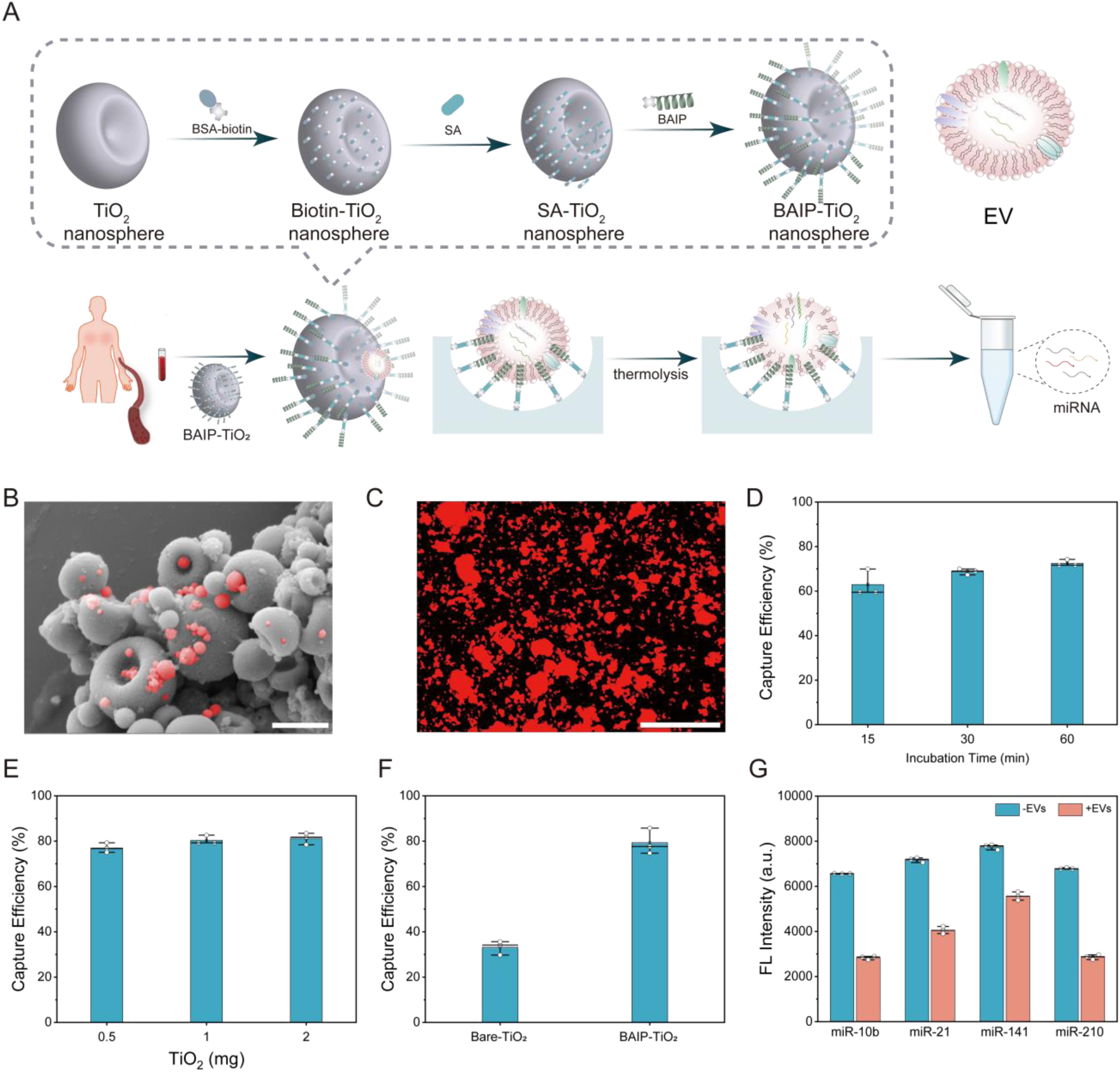
Characterization of isolated EVs and optimization of capture conditions for the BAIP-TiO_2_. (A) Isolation of EVs from serum samples using BAIP-TiO_2_ and releasing target miRNAs via thermal lysis. Insert: Schematic illustration of the surface functionalization of bowl-like TiO_2_ nanospheres with BAIP. (B) SEM image of the captured EVs on BAIP-TiO_2_. Scale bar = 1 μm. (C) Fluorescence images of the captured EVs by BAIP-TiO_2_, labeled with anti-CD9 antibody (Cy3-labeled goat anti-mouse IgG). Scale bar = 100 μm. (D) and (E) Effect of incubation time and BAIP-TiO_2_ dosage on capture efficiency. (F) Comparison of capture efficiency between bare TiO_2_ and BAIP-TiO_2_. Model EVs concentration: 1 ×10^10^ particles /mL. (G) BAIP-TiO_2_-mediated EVs miRNAs detection.

### Detection of EV Derived miRNAs in Clinical Samples

The practical workflow for multiplexed analysis of EVs derived miRNAs in clinical diagnosis was illustrated in **Fig. 6A**. To evaluate the clinical applicability of the **Cap-DISCs** platform, a total of 65 serum samples were collected, including healthy donors (HD, *n* = 20), breast cancer patients (BC, *n* = 26) and triple-negative breast cancer patients (TNBC, *n* = 19). The expression levels of four selected miRNAs were quantified for each sample using established DISCs assay. As shown in **Fig. 6B**, EVs miRNA derived from cancer patients exhibited significantly elevated ΔFL intensity compared with those from healthy donors, indicating markedly elevated expression levels of all four studied miRNA biomarkers in cancer patient-derived EVs (*P* < 0.0001). This result highlights the strong association between these EV encapsulated miRNAs and tumor related pathological states.

Consistently, the heatmap analysis further revealed distinct expression patterns among different groups, with EVs isolated from TNBC patients exhibiting highest overall miRNA abundance, suggesting a more aggressive molecular phenotype (**Fig. 6C**). We next evaluated the diagnostic performance of individual miRNAs and their combined signatures for differentiating among HD, BC and TNBC groups. Receiver operating characteristic (ROC) analysis demonstrated that both single miRNA markers and the multi-miRNA panel exhibited effective diagnostic capability for distinguishing HD from BC, and HD from TNBC (**Fig. S8A** and **Fig. S8B**). Notably, the performance of individual miRNAs was comparable in these binary classifications, indicating that each biomarker contributes meaningful diagnostic information.

More importantly, when distinguishing between BC and TNBC, the multi-miRNA panel exhibited substantially improved discrimination compared with individual markers. As shown in **Fig. 6D**, the area under the curve increased markedly from 0.704 for single miRNAs to 0.972 for the combined panel. This significant enhancement could be attributed to the integration of multiple molecular features, which reduced the impact of biological variability and improved classification robustness. In summary, these results demonstrated that the **DISCs** platform enabled reliable multiplexed detection of EV derived miRNAs in clinical samples and provided enhanced diagnostic accuracy through multi-marker integration, highlighting its potential for precise cancer stratification and clinical decision support.

Machine learning has emerged as a powerful tool for extracting meaningful patterns from complex biological datasets and improving diagnostic accuracy. Within the domain of cancer screening and classification, algorithms such as principal component analysis (PCA), linear discriminant analysis (LDA) and eXtreme Gradient Boosting (XGBoost) were widely used for dimensional reduction and classification, thereby enhancing the performance of predictive models. Motivated by this, we attempted to leverage multiple machine learning approaches to further improve the discrimination between the BC and TNBC based on multiplexed miRNA signatures. PCA was initially performed to visualize the distribution of the EV derived miRNA profiles in the clinical cohort. As shown in **Fig. 6E**, the HD group displayed a well-separated cluster, indicating distinct molecular features compared with cancer samples. However, the BC and TNBC groups exhibited substantial overlap, suggesting that the unsupervised dimensionality reduction alone was insufficient to resolve subtle differences between these subtypes.

To improve classification performance, the LDA was subsequently applied to encode the ΔFL intensity data onto a discriminative subspace. As shown in **Fig. 6F**, the canonical score plot revealed clearer separation among the three groups. The corresponding confusion matrix further quantified the classification performance, yielding an overall accuracy of 89% for distinguishing the TNBC from BC (**Fig. 6G**) and overall accuracy of 98% for distinguishing the BC from HD (**Fig. S8C)**. Although all 23 BC patients and 16 TNBC patients were correctly classified, misclassifications still occurred, indicating that linear models might have limited capability in capturing complex nonlinear relationships within the dataset. To further enhance diagnostic accuracy, an eXtreme Gradient Boosting model was constructed. The dataset of 65 clinical samples was randomly divided into a training set and a test set with a ratio of 8:2. As shown in **Fig. 6H**, the integrated analysis of four miRNAs achieved 100% classification accuracy for distinguishing BC patients from TNBC patients in both training and test cohorts. This performance was markedly superior to that of individual miRNA markers (**Table S3**), highlighting the advantage of multi feature integration combined with nonlinear learning algorithms. These results demonstrated that integrating multiplexed miRNA signatures with advanced machine learning algorithms significantly improved subtype classification and enabled more precise discrimination of TNBC from other BC patients and HD individuals. The improved performance of the XGBoost model could be attributed to its ability to capture complex interactions among multiple biomarkers and to model nonlinear decision boundaries, thereby enhancing both sensitivity and specificity. Collectively, the combination of efficient EVs isolation, sensitive DISCs based detection, and data driven machine learning analysis provided a robust strategy for accurate identification of cancer subtypes. This integrated approach held strong potential for improving the diagnostic performance of EVs-based liquid biopsy in clinical settings.

**Figure 6.**
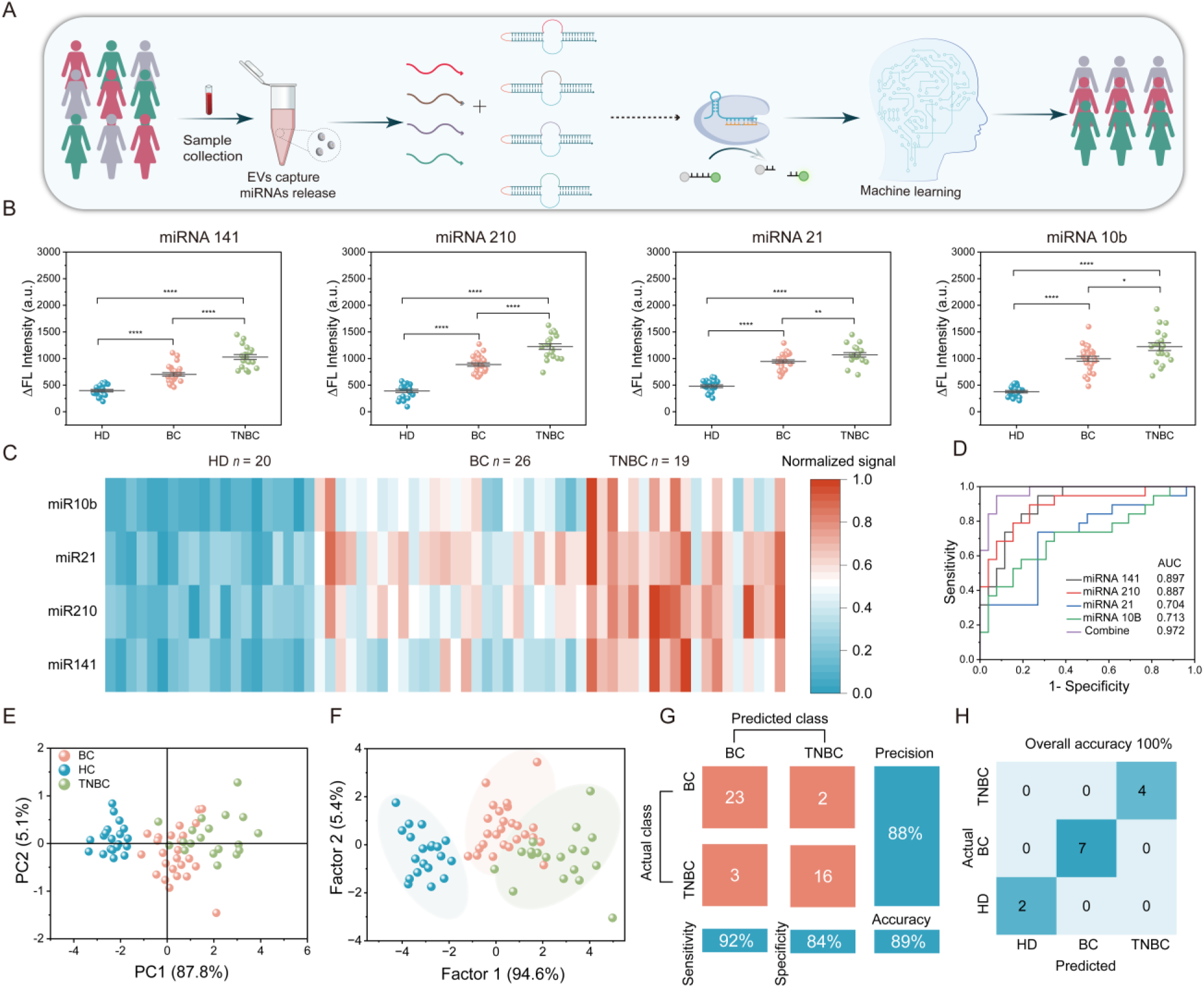
Profiling of isolated EVs miRNA in serum samples to distinguish cancer patients from healthy donors and achieve subtype differentiation. (A) Schematic illustration of the miRNA profiling in the isolated EVs from clinical samples. (B) Scatter plot and (C) heatmap analysis to compare the expression levels of miR-141, miR-210, miR-21, and miR-10b on isolated serum-derived EVs between patients and healthy donors. (D) ROC curves between BC and TNBC, evaluated by expression levels of miR-141, miR-210, miR-21, and miR-10b on isolated serum-derived EVs between BC and TNBC. (E) PCA and (F) LDA distinguished between HD, BC and TNBC using four miRNAs in isolated EVs as the input. (G) Confusion matrix using LDA to differentiate BC and TNBC. (H) Confusion matrix using XGBoost distinguished between HD, BC and TNBC. \**P* <0.05, \*\**P* < 0.01, \*\*\**P* < 0.001, \*\*\*\**P* < 0.0001 indicate significant difference.

## CONCLUSION

In summary, we found that the introduction of ribonucleotides into ssDNA activators could influence the split DNA target-mediated activation of Cas12a. By combining this feature with an enzyme-free amplification strategy based on DNAzyme-mediated target-triggered sequential cleavage mechanism, we achieved sensitive detection of miRNAs derived from EVs. Additionally, to address limitations of effective EV isolation, we developed a dual-recognition strategy for capturing EVs, enabling rapid and highly efficient isolation EVs. By promoting the rapid release of captured EVs miRNAs through thermal lysis, we successfully established an integrated capture-and-detection biosensing platform (**Cap-DISCs**). Furthermore, we evaluated the diagnostic performance of this method for breast cancer (BC) and triple-negative breast cancer (TNBC) subtypes in a clinical cohort (*n* = 65). By integrating four miRNA biomarkers, the biosensor achieved 100% accuracy in distinguishing between breast cancer and TNBC patients in both the training and testing sets. Overall, the established method improves the diagnostic accuracy of cancer and demonstrates strong ability to distinguish cancer subtypes.

## ASSOCIATED CONTENT

### Data Availability Statement

The research data presented in this study are available on request from the corresponding author.

### Supporting Information

The Supporting Information is available free of charge at http://xxxxx/xxxxx.

Material and methods, the sequence of all the used DNAs and RNAs, DNA design strategies and some experimental results, including the feasibility, optimization of reaction condition, the characterization of TiO_2_ and EVs, and so on (DOX)

## ACKNOWLEDGEMENTS

This work is supported by the National Natural Science Foundation of China (Grant Nos. 22174049, 22574059), the Natural Science Foundation of Hubei Province of China (No. 2021CFB335). The authors also thank the Analytical and Testing Center of HUST and the Medical Subcenter of HUST Analytical & Testing Center for material characterization and data acquisition.

## Table of Contents

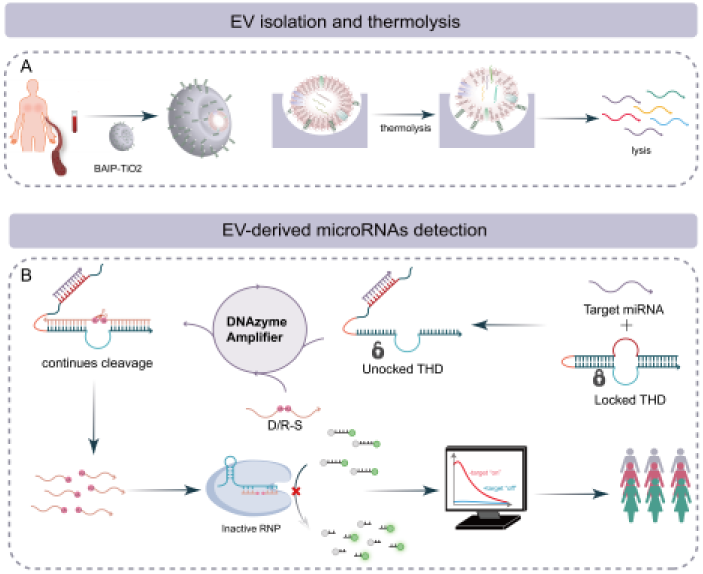

